# Genome-wide association study of depression phenotypes in UK Biobank (n = 322,580) identifies the enrichment of variants in excitatory synaptic pathways

**DOI:** 10.1101/168732

**Authors:** David M. Howard, Mark J. Adams, Masoud Shirali, Toni-Kim Clarke, Riccardo E. Marioni, Gail Davies, Jonathan R. I. Coleman, Clara Alloza, Xueyi Shen, Miruna C. Barbu, Eleanor M. Wigmore, Jude Gibson, Saskia P. Hagenaars, Cathryn M. Lewis, Daniel J. Smith, Patrick F. Sullivan, Chris S. Haley, Gerome Breen, Ian J. Deary, Andrew M. McIntosh

**Author notes:** Corresponding author: David M. Howard Division of Psychiatry, University of Edinburgh, Royal Edinburgh Hospital, Edinburgh, UK.

## Abstract

Depression is a polygenic trait that causes extensive periods of disability and increases the risk of suicide, a leading cause of death in young people. Previous genetic studies have identified a number of common risk variants which have increased in number in line with increasing sample sizes. We conducted a genome-wide association study (GWAS) in the largest single population-based cohort to date, UK Biobank. This allowed us to estimate the effects of ≈ 8 million genetic variants in 320,000 people for three depression phenotypes: broad depression, probable major depressive disorder (MDD), and International Classification of Diseases (ICD, version 9 or 10)-coded MDD. Each phenotype was found to be significantly genetically correlated with the results from a previous independent study of clinically defined MDD. We identified 14 independent loci that were significantly associated (*P* < 5 × 10^−8^) with broad depression, two independent variants for probable MDD, and one independent variant for ICD-coded MDD. Gene-based analysis of our GWAS results with MAGMA revealed 46 regions significantly associated (*P* < 2.77 × 10^−6^) with broad depression, two significant regions for probable MDD and one significant region for ICD-coded MDD. Gene region-based analysis of our GWAS results with MAGMA revealed 59 regions significantly associated (*P* < 6.02 × 10^−6^) with broad depression, of which 27 were also detected by gene-based analysis. Variants for broad depression were enriched in pathways for excitatory neurotransmission, mechanosensory behavior, postsynapse, neuron spine and dendrite. This study provides a number of novel genetic risk variants that can be leveraged to elucidate the mechanisms of MDD and low mood.

## Introduction

Depression is ranked as the largest contributor to global disability affecting 322 million people^1^. The heritability (h^2^) of major depressive disorder (MDD) is estimated at 37% from twin studies^2^ and common single nucleotide polymorphisms (SNPs) contribute approximately 9% to variation in liability^3^, providing strong evidence of a genetic contribution to its causation. Previous genetic association studies have used a number of depression phenotypes, including self-declared depression^4^, clinician diagnosed MDD^5^ and depression ascertained via hospital records^6^, with some evidence of overlapping genetic architecture between a subset of these definitions. Different definitions of depression are rarely included in large sample studies, although UK Biobank is an exception. The favouring of greater sample size over clinical precision has yielded a steady increase over time in the number of variants for ever more diverse MDD phenotypes^3-5,7^. In the current paper, we extend this approach to the study of three depression-related phenotypes within the large UK Biobank cohort and identify new disease biology based upon our findings.

The UK Biobank cohort provides data on over 500,000 individuals and represents an opportunity to conduct the largest genome-wide association study (GWAS) of depression to date within a single cohort. This cohort has been extensively phenotyped allowing us to derive three depression traits: self-reported past help-seeking for problems with ‘nerves, anxiety, tension or depression’ (hereby termed ‘broad depression’); self-reported depressive symptoms with associated impairment (termed ‘probable MDD’); and MDD identified from International Classification of Diseases (ICD)-9 or ICD-10 hospital admission records (termed ICD-coded MDD). We also conducted a gene-based analyses with the MAGMA software package^8^ to identify genes, regions and pathways associated with each phenotype and used GTEx^9^ to identify if the significant variants identified were expression quantitative trait loci (eQTL).

## Materials and Methods

The UK Biobank cohort is a population-based cohort consisting of 501,726 individuals, recruited at 23 centres across the United Kingdom. Genotypic data was available for 488,380 individuals and was imputed with IMPUTE4 and used the HRC reference panel^10^ to identify ≈ 19M variants for 487,409 individuals^11^. We excluded 131,790 related individuals based on a shared relatedness of up to the third degree using kinship coefficients (> 0.044) calculated using the KING toolset^12^, and excluded a further 79,990 individuals that were either not recorded as “white British”, outliers based on heterozygosity, or had a variant call rate < 98%. We subsequently added back in one member of each group of related individuals by creating a genomic relationship matrix and selected individuals with a genetic relatedness less than 0.025 with any other participant (n = 55,745). We removed variants with a call rate < 98%, a minor allele frequency < 0.01, those that deviated from Hardy-Weinberg equilibrium (*P* < 10^−6^), or had an imputation accuracy score < 0.1 leaving a total of 7,826,341 variants for 331,374 individuals.

Extensive phenotypic data were collected for UK Biobank participants using health records, biological sampling, physical measures, and touchscreen tests and questionnaires. We used three definitions of depression in the UK Biobank sample, which are explained in greater depth in the Supplementary Information and are summarised below.

### *Broad* depression *phenotype*

The broadest phenotype (broad depression) was defined using self-reported help-seeking behaviour for mental health difficulties. Case and control status was determined by the touchscreen response to either of two questions ‘Have you ever seen a general practitioner (GP) for nerves, anxiety, tension or depression?’ or ‘Have you ever seen a psychiatrist for nerves, anxiety, tension or depression?. Caseness for broad depression was determined by answering ‘Yes’ to either question at either the initial assessment visit or at any repeat assessment visit or if there was a primary or secondary diagnosis of a depressive mood disorder from linked hospital admission records. The remaining respondents were classed as controls if they provided ‘No’ responses to both questions during all assessments that they participated in.

### Probable MDD phenotype

The second depression phenotype (probable MDD) was derived from touchscreen responses to questions about the presence and duration of low mood and anhedonia, following the definitions from Smith, et al. ^13^, whereby the participant had indicated that they were ‘Depressed/down for a whole week; plus at least two weeks duration; plus ever seen a GP or psychiatrist for ‘nerves, anxiety, or depression’ OR ever anhedonia for a whole week; plus at least two weeks duration; plus ever seen a GP or psychiatrist for ‘nerves, anxiety, or depression’. Cases for the probable MDD definition were supplemented by diagnoses of depressive mood disorder from linked hospital admission records.

### ICD-coded phenotype

The ICD-coded MDD phenotype was derived from linked hospital admission records. Participants were classified as cases if they had either an ICD-10 primary or secondary diagnosis for a mood disorder. ICD-coded MDD controls were participants who had linked hospital records, but who did not have any diagnosis of a mood disorder and were not probable MDD cases.

For the three UK Biobank depression phenotypes we excluded: participants who were identified with bipolar disorder, schizophrenia, or personality disorder using self-declared data, touchscreen responses (per Smith, et al. ^13^), or ICD codes from hospital admission records; and participants who reported having a prescription for an antipsychotic medication during a verbal interview. Further exclusions were applied to control individuals if they had a diagnosis of a depressive mood disorder from hospital admission records, had reported having a prescription for antidepressants, or self-reported depression (see Supplementary Information for full phenotype criteria and UK Biobank field codes). This provided a total of 113,769 cases and 208,811 controls (n_total_ = 322,580, prevalence = 35.27%) for the broad depression phenotype, 30,603 cases and 143,916 controls (n_total_ = 174,519, prevalence = 17.54%) for the probable MDD phenotype, and 8,276 cases and 209,308 controls (n_total_ = 217,584, prevalence = 3.80%) for the ICD-coded MDD phenotype.

To validate the three phenotypes we derived for the UK Biobank cohort, genetic correlations were calculated using Linkage Disequilibrium Score regression (LDSR)^14^ using summary statistics from the Major Depressive Disorder Working Group of the Psychiatric Genomics Consortium. ^5^ study that used a clinically derived phenotype for MDD. We also calculated the genetic correlation with a neuroticism phenotype^15^.

### Association analysis

We performed a linear association test to assess the effect of each variant using BGENIE v1.1^11^:

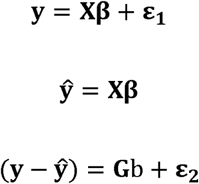

where **y** was the vector of binary observations for each phenotype (controls coded as 0 and cases coded as 1). **β** was the matrix of fixed effects, including sex, age, genotyping array, and 8 principal components and **X** was the corresponding incidence matrices. **(y – ŷ)** was a vector of phenotypes residualized on the fixed effect predictors, **G** was a vector of expected genotype counts of the effect allele (dosages), b was the effect of the genotype on residualized phenotypes, and **ε**_1_ and **ε**_2_ were vectors of normally distributed errors.

Genome-wide statistical significance was determined by the conventional threshold of a *P*-value of association < 5 × 10^−8^. To determine significant variants that were independent the clump command in Plink 1.90b4^16^ was applied using --clump-p1 1e-4 --clump-p2 1e-4 --clump-r2 0.1 --clump-kb 3000, mirroring the approach of Major Depressive Disorder Working Group of the Psychiatric Genomics Consortium., et al. ^3^. Therefore variants which were within 3Mb of each other and shared a linkage disequilibrium greater than 0.1 were clumped together and only the most significant variant reported. Due to the complexity of major histocompatibility complex (MHC) region an approach similar to that of The Schizophrenia Psychiatric Genome-Wide Association Study ^17^ was taken and only the most significant variant across that region is reported.

LDSR^14^ was used to provide a SNP-based estimate of the heritability of the phenotypes using the whole-genome summary statistics obtained by the association analyses. LDSR was also used to examine the data for evidence of inflation of the test statistics, based on the intercept, due to population stratification.

### Gene- and region-based analyses

Two downstream analyses of the results were conducted using MAGMA^8^ (Multi-marker Analysis of GenoMic Annotation) by applying a principal component regression model to the results of our association analyses. In the first downstream analysis, a gene-based analysis was performed for each phenotype using the results from our GWAS. Genetic variants were assigned to genes based on their position according to the NCBI 37.3 build, resulting in a total of 18,033 genes being analysed. The European panel of the 1,000 Genomes data (phase 1, release 3)^18^ was used as a reference panel to account for linkage disequilibrium. A genome-wide significance threshold for gene-based associations was calculated using the Bonferroni method (α = 0.05 / 18,033; *P* < 2.77 × 10^−6^).

In the second downstream analysis, a region-based analysis was performed for each phenotype. To determine the regions, haplotype blocks identified by recombination hotspots were used as described by Shirali, et al. ^19^ and implemented in an analysis of MDD by Zeng, et al. ^20^ for detecting causal regions. Block boundaries were defined by hotspots of at least 30 cM per Mb based on a European subset of the 1,000 genome project recombination rates. This resulted in a total of 8,308 regions being analysed using the European panel of the 1,000 Genomes data (phase 1, release 3)^18^ as a reference panel to account for linkage disequilibrium. A genome-wide significance threshold for region-based associations was calculated using the Bonferroni correction method (α = 0.05 / 8,308; *P* < 6.02 × 10^−6^).

### Pathway analysis

The pathway analysis was performed on our gene-based analysis results. The analysis was a gene-set enrichment analysis that was conducted utilising gene-annotation files from the Gene Ontology (GO) Consortium (http://geneontology.org/)^21^ taken from the Molecular Signatures Database (MSigDB) v5.2^22^. The GO consortium includes gene-sets for three ontologies; molecular function, cellular components and biological function. This annotation file consisted of 5,917 gene-sets which were corrected for multiple testing correction using the MAGMA default setting correcting for 10,000 permutations. Visualisation of pathways was obtained using the online tool, GeneMANIA^23^.

### eQTL identification

The online GTEx portal (https://www.gtexportal.org/home/) was used to determine whether any of the genome-wide significant variants for each phenotype were eQTL^9^.

## Results

We conducted a genome-wide association study testing the effect of 7,826,341 variants on three depression phenotypes using up to 322,580 UK Biobank participants. The study demographics for each UK Biobank phenotype and within the case and control groups are provided in Table 1.

**Table 1.** Number of individuals, number of each sex, mean age in years, age range in years for each of the assessed UK Biobank phenotypes and within the respective case and control groups

| Phenotype | Status | N | Males | Females | Mean Age (st.dev) | Age Range |
| --- | --- | --- | --- | --- | --- | --- |
| Broad depression | Cases | 113,769 | 40,477 | 73,292 | 56.5 (7.8) | 39-73 |
|  | Controls | 208,811 | 109,426 | 99,385 | 57.1 (8.1) | 39-72 |
|  | Total | 322,580 | 149,903 | 172,677 | 56.9 (8.0) | 39-73 |
| Probable MDD | Cases | 30,603 | 11,346 | 19,257 | 56.1 (7.8) | 40-70 |
|  | Controls | 143,916 | 65,015 | 78,901 | 57.1 (7.9) | 39-73 |
|  | Total | 174,519 | 76,361 | 98,158 | 56.9 (7.9) | 39-73 |
| ICD-coded MDD | Cases | 8,276 | 3,098 | 5,178 | 56.5 (7.9) | 40-70 |
|  | Controls | 209,308 | 99,961 | 109,347 | 57.6 (8.0) | 39-73 |
|  | Total | 217,584 | 103,059 | 114,525 | 57.6 (8.0) | 39-73 |

The estimated SNP-based heritabilities, genetic correlations between each UK Biobank phenotype and genetic correlations with a clinically defined MDD phenotype and obtained from the study conducted by the Major Depressive Disorder Working Group of the Psychiatric Genomics Consortium. ^5^ and a neuroticism phenotype^15^ for each UK Biobank phenotype are provided in Table 2.

**Table 2.** The SNP-based heritability (h^2^), the genetic correlations (r_g_) between each UK Biobank phenotype and the r_g_ with a depression and a neuroticism phenotype obtained from separate studies† for each of the assessed UK Biobank phenotypes.

| Phenotype | $h^2$ (s.e) | $r_g$ with MDD<sup>†</sup> (s.e.) | $r_g$ with neuroticism<sup>†</sup> (s.e.) | $r_g$ with broad depression (s.e.) | $r_g$ with probable MDD (s.e.) |
| --- | --- | --- | --- | --- | --- |
| Broad depression | 0.102 (0.004) | 0.804 (0.092) | 0.671 (0.018) | - | 0.871 (0.050) |
| Probable MDD | 0.047 (0.006) | 0.695 (0.139) | 0.518 (0.050) | 0.871 (0.050) | - |
| ICD-coded MDD | 0.100 (0.012) | 0.669 (0.122) | 0.563 (0.045) | 0.863 (0.046) | 0.848 (0.052) |

<sup>†</sup>To conduct the genetic correlations with the UK Biobank phenotypes the depression phenotype from Major Depressive Disorder Working Group of the Psychiatric Genomics Consortium.<sup>5</sup> and the neuroticism phenotype<sup>15</sup> was used

There were 1,643 variants that were genome-wide significant (*P* < 5 × 10^−8^) for an association with broad depression, of which 14 were independent (Table 3). The association analysis of probable MDD identified 20 variants with *P* < 5 × 10^−8^ and of these two were independent (Table 4). There was one independent genome-wide significant variant for ICD-coded MDD (Table 5). Manhattan plots of all the variants analysed are provided in Figures 1, 2, and 3 for broad depression, probable MDD, and ICD-coded MDD, respectively. Q-Q plots of the observed *P*-values on those expected are provided in Supplementary Figures 1, 2, and 3 for broad depression, probable MDD, and ICD-coded MDD, respectively. There were 4,390, 189 and 108 variants with *P* < 1 × 10^−6^ for an association with broad depression (see Supplementary Table 1), probable MDD (see Supplementary Table 2), and ICD-coded MDD (see Supplementary Table 3), respectively. None of the phenotypes examined provided evidence of inflation of the test statistics due to population stratification (see Supplementary Table 4).

**Table 3.** Independent variants with a genome-wide significant (*P* < 5 × 10^−8^) association with broad depression

| Chr | Marker name | Position | A1 | A2 | Allele Frequency | Imputation Accuracy | Beta | Standard error | Gene +/- 10kb | $P$-value |
| --- | --- | --- | --- | --- | --- | --- | --- | --- | --- | --- |
| 1 | rs10127497 | 67050144 | T | A | 0.144 | 1.00 | 0.0048 | 0.0009 | *SGIP1* | $1.29 \times 10^{-8}$ |
| 1 | rs6699744 | 72825144 | T | A | 0.617 | 1.00 | 0.0061 | 0.0008 | - | $1.64 \times 10^{-13}$ |
| 1 | rs6424532 | 73664022 | A | G | 0.478 | 1.00 | 0.0046 | 0.0008 | - | $3.91 \times 10^{-8}$ |
| 1 | rs7548151 | 177026983 | A | G | 0.089 | 1.00 | 0.0051 | 0.0009 | *ASTN1* | $3.87 \times 10^{-9}$ |
| 5 | rs40465 | 103981726 | G | T | 0.325 | 1.00 | 0.0052 | 0.0008 | *RP11-6N13.1* | $4.45 \times 10^{-10}$ |
| 6 | rs3094054 | 30333505 | T | G | 0.136 | 1.00 | -0.0061 | 0.0008 | - | $1.79 \times 10^{-13}$ |
| 6 | rs112348907 | 73587953 | G | A | 0.292 | 1.00 | 0.0047 | 0.0008 | | $1.52 \times 10^{-8}$ |
| 7 | rs3807865 | 12250402 | A | G | 0.421 | 1.00 | 0.0057 | 0.0008 | *TMEM106B* | $7.28 \times 10^{-12}$ |
| 7 | rs2402273 | 117600424 | C | T | 0.405 | 1.00 | 0.0050 | 0.0008 | - | $1.95 \times 10^{-9}$ |
| 9 | rs263575 | 17033840 | A | G | 0.466 | 1.00 | -0.0047 | 0.0008 | - | $2.31 \times 10^{-8}$ |
| 10 | rs1021363 | 106610839 | G | A | 0.648 | 1.00 | -0.0048 | 0.0008 | *SORCS3* | $1.04 \times 10^{-8}$ |
| 11 | rs10501696 | 88748162 | G | A | 0.495 | 0.99 | -0.0055 | 0.0008 | *GRM5* | $6.73 \times 10^{-11}$ |
| 13 | rs9530139 | 31847324 | T | C | 0.195 | 1.00 | -0.0049 | 0.0008 | *B3GLCT* | $2.63 \times 10^{-9}$ |
| 15 | rs28541419 | 88945878 | G | C | 0.231 | 1.00 | -0.0046 | 0.0008 | - | $2.78 \times 10^{-8}$ |

The allele frequency and reported effect size (beta) is for the A1 allele. The chromosome (Chr) and position in Mb is given with regards to the GRCh37 assembly.

**Table 4.** Independent variants with a genome-wide significant (*P* < 5 × 10^−8^) association with probable MDD

| Chr | Marker name | Position | A1 | A2 | Allele Frequency | Imputation Accuracy | Beta | Standard error | Gene +/- 10kb | P-value |
| --- | --- | --- | --- | --- | --- | --- | --- | --- | --- | --- |
| 2 | rs10929355 | 15398964 | G | T | 0.470 | 1.00 | -0.0053 | 0.0009 | *NBAS* | $5.84 \times 10^{-9}$ |
| 7 | rs5011432 | 12268668 | C | A | 0.418 | 1.00 | 0.0051 | 0.0009 | *TMEM106B* | $2.23 \times 10^{-8}$ |

The allele frequency and reported effect size (beta) is for the A1 allele. The chromosome (Chr) and position in Mb is given with regards to the GRCh37 assembly.

**Table 5.** Independent variants with a genome-wide significant (*P* < 5 × 10^−8^) association with ICD-coded MDD

| Chr | Marker name | Position | A1 | A2 | Allele Frequency | Imputation Accuracy | Beta | Standard error | Gene +/- 10kb | P-value |
| --- | --- | --- | --- | --- | --- | --- | --- | --- | --- | --- |
| 7 | rs1554505 | 1983929 | A | G | 0.742 | 1.00 | 0.0025 | 0.0004 | *MAD1L1* | $2.74 \times 10^{-9}$ |

The allele frequency and reported effect size (beta) is for the A1 allele. The chromosome (Chr) and position in Mb is given with regards to the GRCh37 assembly.

**Figure 1.**
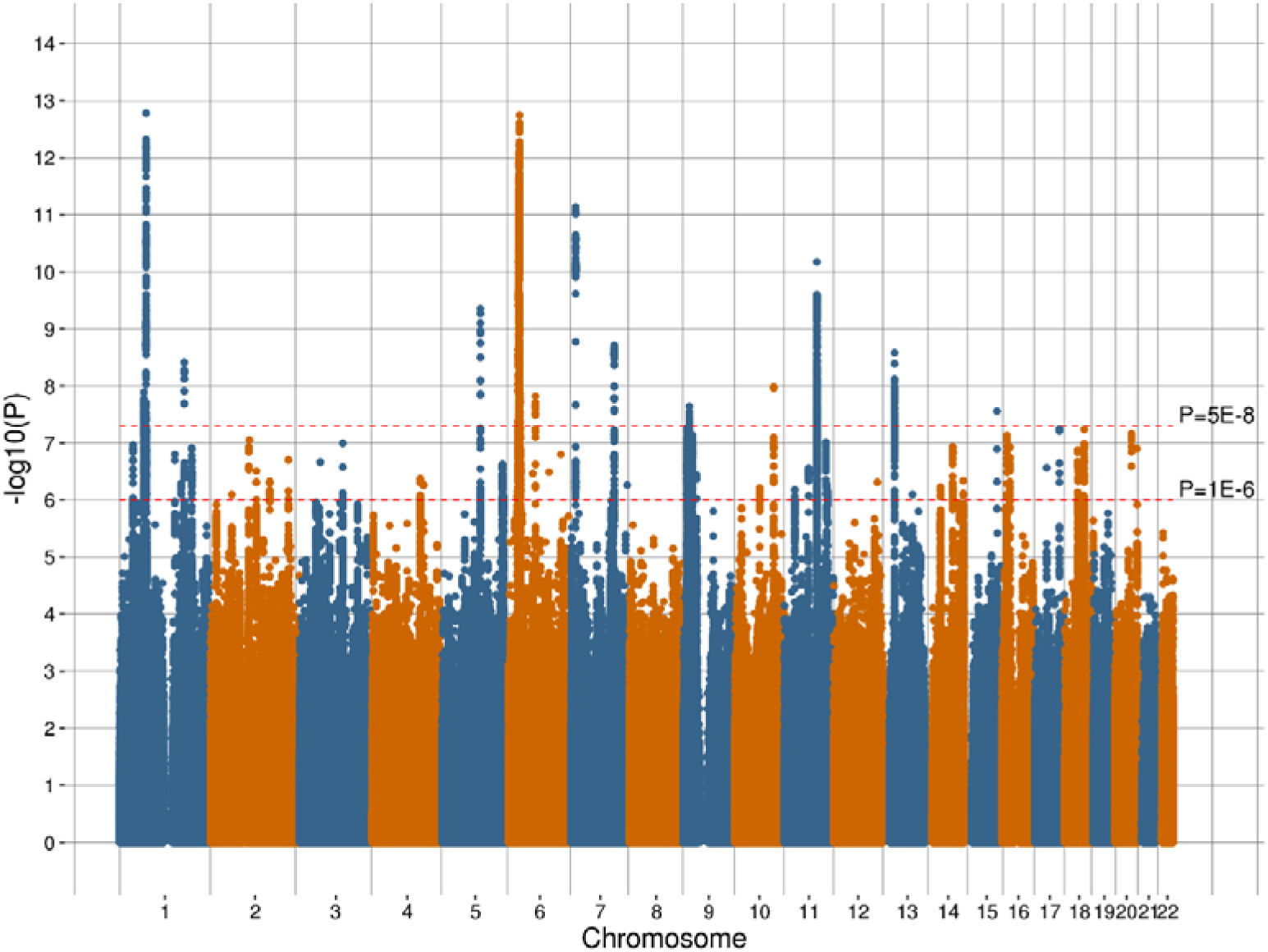
Manhattan plot of the observed –log_10_ *P*-values of each variant for an association with broad depression in the UK Biobank cohort. Variants are positioned according to the GRCh37 assembly.

**Figure 2.**
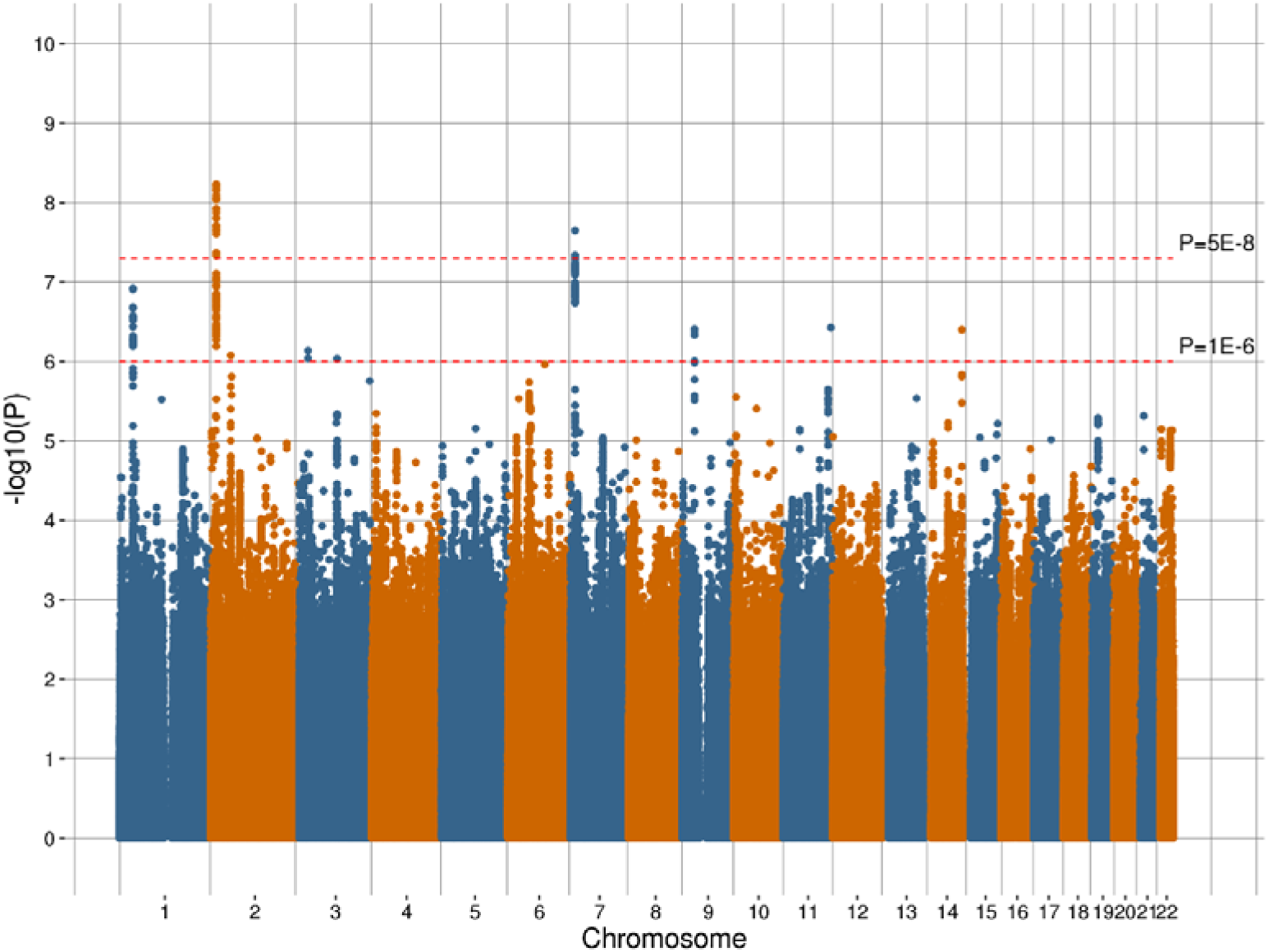
Manhattan plot of the observed –log_10_ *P*-values of each variant for an association with probable MDD in the UK Biobank cohort. Variants are positioned according to the GRCh37 assembly.

**Figure 3.**
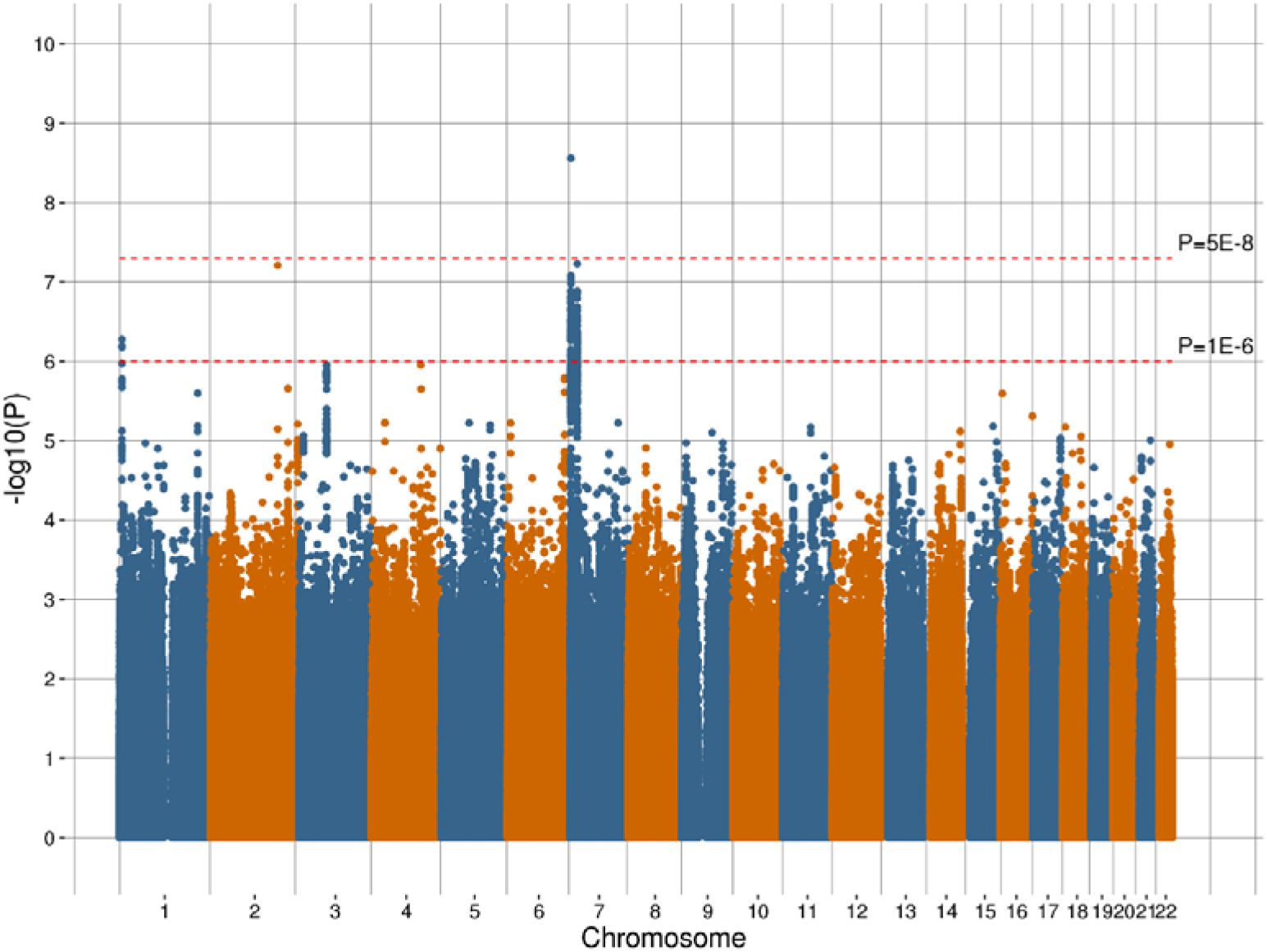
Manhattan plot of the observed –log_10_ *P*-values of each variant for an association with ICD-coded MDD in the UK Biobank cohort. Variants are positioned according to the GRCh37 assembly.

### Gene- and region-based analyses

The MAGMA package was used to identify gene-based regions with a significant effect (P < 2.77 × 10^−6^) on each trait. This gene-based approach detected 46 significant regions containing 78 genes that were significantly associated with broad depression (Supplementary Table 5), two significant regions containing two genes that were significantly associated with probable MDD (Supplementary Table 6), and one significant region containing one gene that was significantly associated with ICD-coded MDD (Supplementary Table 7).

MAGMA was also used to identify genomic regions, defined by recombination hotspots, with a statistically significant effect (*P* < 6.02 × 10^−6^) on each phenotype. There were 59 significant regions identified for broad depression and a further four regions identified for probable MDD and four regions for ICD-coded MDD. Further details regarding these regions are provided in Supplementary Tables 8, 9, and 10 for broad depression, probable MDD, and ICD-coded MDD, respectively.

Manhattan plots of the gene and regions analysed are provided in Supplementary Figures 4, 5, and 6 for broad depression, probable MDD, and ICD-coded MDD, respectively.

### Pathway analysis

Gene-set enrichment analysis identified five significant pathways for broad depression after applying correction for multiple testing; excitatory synapse (*P*_corrected_ = 0.004, beta = 0.346, s.e. = 0.069), mechanosensory behavior (*P*_corrected_ = 0.009, beta = 1.390, s.e. = 0.290), postsynapse (*P*_corrected_ = 0.009, beta = 0.241, s.e. = 0.050), neuron spine (*P*_corrected_ = 0.035, beta = 0.376, s.e. = 0.085) and dendrite (*P*_corrected_ = 0.041, beta = 0.195, s.e. = 0.045) (Table 6). Figure 4 illustrates the genes and shared protein domains involved in the mechanosensory behaviour pathway. No significant pathways (*P* > 0.05) were associated with probable MDD or ICD-coded MDD after multiple testing correction.

**Table 6.** Pathways with a significant effect (*P*_corrected_ < 0.05) on broad depression following multiple testing correction identified through gene-set enrichment analysis

| Phenotype | Pathway | Number of genes | Beta | Standard error | P-value | $P_{\text{Corrected}}$ |
| --- | --- | --- | --- | --- | --- | --- |
| Broad depression | Excitatory Synapse | 182 | 0.346 | 0.069 | $2.38 \times 10^{-7}$ | 0.004 |
| | Mechanosensory Behavior | 11 | 1.390 | 0.290 | $7.99 \times 10^{-7}$ | 0.009 |
| | Postsynapse | 352 | 0.241 | 0.050 | $8.26 \times 10^{-7}$ | 0.009 |
| | Neuron Spine | 114 | 0.376 | 0.085 | $4.89 \times 10^{-6}$ | 0.035 |
| | Dendrite | 423 | 0.195 | 0.045 | $6.06 \times 10^{-6}$ | 0.041 |

**Figure 4.**
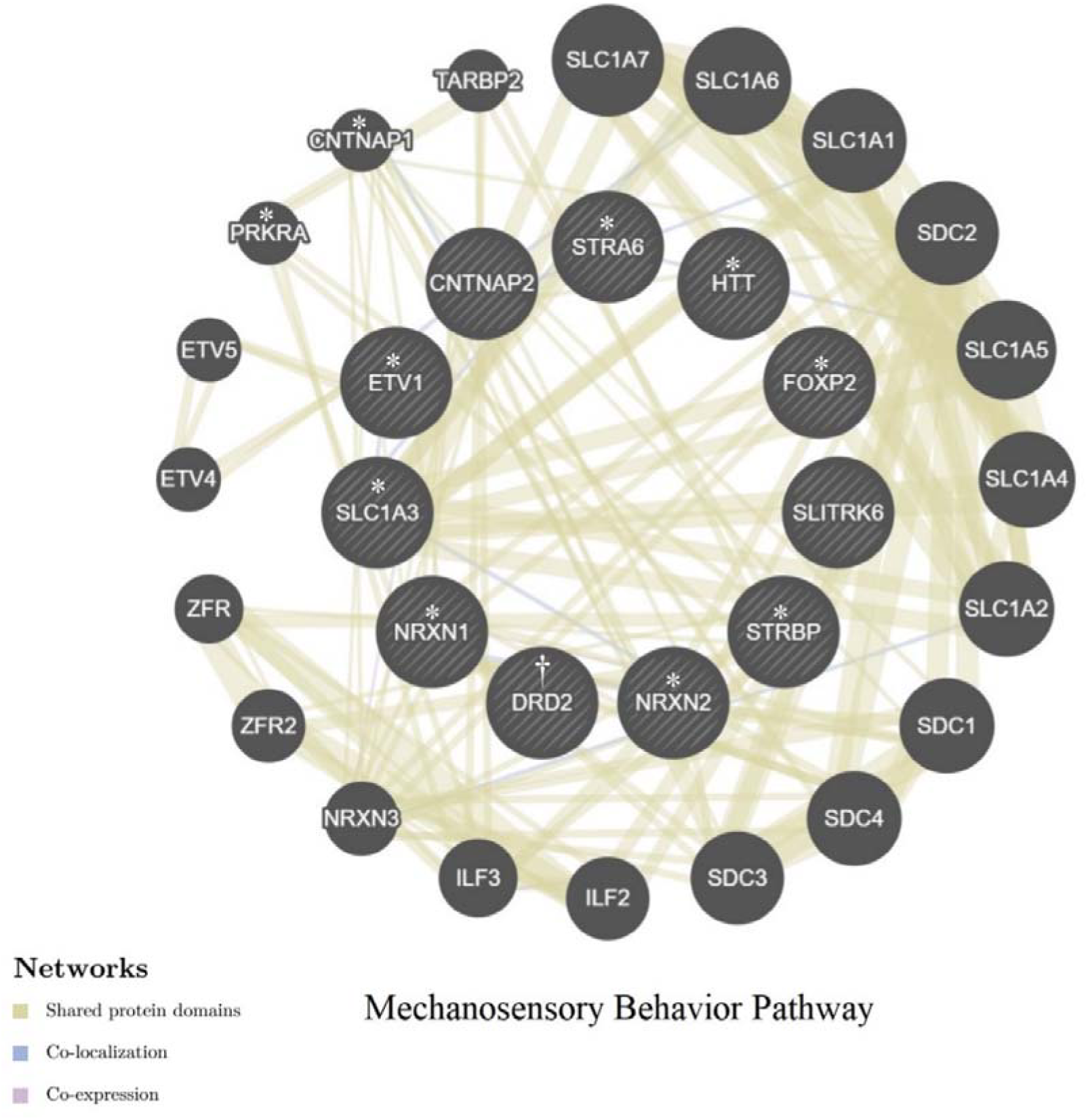
Plot of the mechanosensory behaviour pathway which was significant associated (*P*_corrected_ = 0.009) with broad depression after multiple testing correction. The nodes represent the gene coding regions within the network with connections between those regions represented as edges. † indicates a gene coding region that was itself significantly associated (*P*_corrected_ < 0.05) with broad depression after multiple testing. * indicates a gene coding region within the pathway that was nominally significantly associated (*P* < 0.05) with broad depression. The size of each node is determined by the effect size of the gene coding region within the pathway based on previous studies.

### eQTL identification

Across the three phenotypes examined seven variants were identified as potential eQTLs (Supplementary Table 11), of these included 4 variants found to be eQTL for brain expressed genes. rs6699744 is associated with the expression of *RPL31P12* in the cerebellum and rs9530139 is associated with expression of *B3GALTL* in the cortex. rs40465 and rs68141011 are broad eQTLs affecting the expression of a number of zinc finger protein encoding genes across various brain tissues including *ZNF391, ZNF204P, ZNF192P1, ZSCAN31* and *ZSCAN23.*

## Discussion

This study describes the largest analysis of depression using a single population-based cohort to date. Up to 322,580 individuals from the UK Biobank cohort were used to test the effect of approximately 8 million genetic variants on three depression phenotypes. A total of 17 independent genome-wide significant (*P* < 5 × 10^−8^) variants were identified across the three phenotypes. The broadest definition of the phenotype, broad depression, providing the greatest number of individuals for analysis and also the largest number of significant hits (14 independent variants). The probable MDD phenotype was obtained using the approach of Smith, et al. ^13^, using touchscreen items that were only administered in the last two years of UK Biobank recruitment, and thus only available for only a portion of the sample, yielded two independent genome-wide significant variants. The strictest phenotype was ICD-based MDD, which was dependent on linked hospital admission records for a primary or secondary diagnosis of a mood disorder, had one independent significant variant.

The three UK Biobank phenotypes for depression all had significant genetic correlations (r_g_ ≥ 0.669 (0.122), *P* ≤ 6.13 × 10^−7^) with the results from a mega-analysis of MDD^5^, based on an anchor set of clinically defined cases. Interestingly, it was the broad depression phenotype that had the highest genetic correlation with that clinically-defined MDD phenotype. Neuroticism and MDD do share similar symptoms and we did identify significant genetic correlations (*P* ≤ 2.77 × 10^−25^) between these phenotypes, although as expected the genetic correlations were not as high as those with clinically diagnosed MDD. Each UK Biobank phenotype produced different variants and genes that demonstrated an association. This variability in the variants underlying each phenotype indicates that depression phenotypes may differ markedly in their tractability for genetic studies, and that they may also have somewhat different aetiologies.

A genome-wide significant variant associated with broad depression and identified as an eQTL by GTEx on chromosome 1, rs6699744 (72,825,144 Mb; *P* = 1.64 × 10^−13^) was close to another significant variant (rs11209948; *P* = 8.38 × 10^−11^) at 72,811,904 Mb associated with MDD within the Hyde, et al. ^4^ study. Both these variants were close to the Neural Growth Regulator 1 (*NEGR1*) gene which was associated with MDD in the Major Depressive Disorder Working Group of the Psychiatric Genomics Consortium., et al. ^3^ study. Another significant variant (rs7548151) for broad depression on chromosome 1, located at 177,026,983 Mb, was found within 30 Kb of the microRNA 488 (*MIR488*) coding region. *MIR488* is a brain-enriched miRNA that has been implicated in post-transcriptional regulation of gene expression affecting both stability and translation of mRNAs. Transcriptome analysis has demonstrated that altered expression of *MIR488* is nominally associated with stress response and panic disorder^24^.

The most proximal gene coding region to the significant variant (rs40465) for broad depression on chromosome 5 was RNU6-334P, which is a pseudogene. Although this variant is located within a region that contains no protein-coding sequences and has no known biological function, the region has been associated with depression and depressive symptoms^7,25^. The GTEx analysis identified this variant as an eQTL for brain expressed genes.

A significant variant (rs1021363, 106,610,839 Mb, *P* = 1.04 × 10^−8^) on chromosome 10 was associated with broad depression in our study and was within 4 Kb of another variant (rs10786831; 106,614,571 Mb; *P* = 8.11 × 10^−9^) found to be associated within the Hyde, et al. ^4^ MDD study and close to a variant (rs61867293, 106,563,924 Mb, *P* = 7.0 × 10^−10^) associated with MDD in the Major Depressive Disorder Working Group of the Psychiatric Genomics Consortium., et al. ^3^ study.

There were up to eight variants in the gene-rich MHC region that could have been classified as independent however the complexity of the genetic architecture across this region may confounded this. Therefore we report only the most significant variant (rs3094054). The MHC region has been associated with both schizophrenia and bipolar disorder across multiple studies^26-28^, as well as an early-onset and recurrent form of depression^29^. A closer examination of the MHC region is certainly warranted with regarding psychiatric disease based on previous studies and the results obtained in this paper.

There was a genome-wide significant variant located on chromosome 11 that overlapped the glutamate metabotropic receptor 5 (*GRM5*) protein coding gene. *GRM5* is expressed in the brain and facilitates glutamatergic neurotransmission. *GRM5* has previously been associated with a range of behavioural and neurological phenotypes such as depression^30^, OCD^31^, epilepsy^32^, smoking^33^, Alzheimer’s disease^34,35^, autism^36^ and schizophrenia^37^. A recent study found a role for Metabotropic glutamate receptor 5 (*mGluR5*) in relation to stress-induced depression in mice^38^ and *GRM5* antagonists have been shown to have anxiolytic and anti-depressant properties^39,40^.

A variant on chromosome 7, rs1554505, was associated with ICD-coded MDD. This variant is located in the MAD1 mitotic arrest deficient-like 1 (*MAD1L1*) gene coding region. *MAD1L1* is a known susceptibility locus for schizophrenia^41-43^ and a recent study showed that differential reward processing during an fMRI task in carriers of a *MAD1L1* bipolar risk allele ^44^.

### Gene-based and region-based analyses

The gene-based analysis identified a total of 79 genome-wide significant (P < 2.77 × 10^−6^) genes across the three phenotypes. The transmembrane protein 106B (*TMEM106B*) gene coding region was identified in both the broad depression and probable MDD. *TMEM106B* encodes a type II transmembrane protein of unknown function, although represents a risk factor for Frontotemporal lobar degeneration, especially in patients with *progranulin* mutations^45^. An inverse relationship between *TMEM106B* (downregulation) and *progranulin* (upregulation) expression levels in Alzheimer’s disease brains has been reported which demonstrated the role of *TMEM106B* gene in the pathological processes of Alzheimer’s disease^46^.

Our region-based analysis identified 59 genome-wide significant (P < 6.02 × 10^−6^) regions across the three phenotypes, with three of these regions detected in more than one phenotype. The region-based method detected regions harbouring several known genes that are reported to have effect on depression and other mental diseases that were not detected in our gene-based analysis.

The region-based analysis of both broad depression and probable MDD detected a significant region on chromosome 1 that contained the glutamate ionotropic receptor kainate type subunit 3 (*GRIK3*) protein coding region. Glutamate receptors are the main excitatory neurotransmitter receptors in the mammalian brain. Moreover, these receptors are active in several neurophysiologic processes. *GRIK3* has been associated with schizophrenia^47,48^, neuroticism^49^ and recurrent MDD^50,51^. Higher levels of *GRIK3* have been reported in MDD suicides compared to MDD non-suicides with *GRIK3* expression a strong predictor of suicide^52^.

The analysis of broad depression and ICD-coded MDD both detected a significant region containing the Receptor tyrosine-protein kinase erbB-4X (*ERBB4*) protein coding region. *ERBB4* is a member of the Tyr protein kinase family and the epidermal growth factor receptor subfamily. *ERBB4* has been previously linked to schizophrenia^53^ and impairments in the link between Neuregulin 1 (*NRG1*) and *ERBB4* signalling are associated with schizophrenia ^54^ and anxiety behaviours^55^.

The broad depression analysis identified a region containing the dihydropyrimidine dehydrogenase (*DPYD*) gene coding region which has been associated with schizophrenia and bipolar disorder^56^ and borderline personality disorder^57^. Also identified were regions containing the Neurexin 1 (*NRXN1*) gene coding region which has been associated with Tourette syndrome^58^ and non-syndromic autism spectrum disorder^59^ and the regulator of G protein signalling 6 (*RGS6*) coding region which has been previously associated with alcoholism^60^, depression/anxiety^61^ and Parkinson’s disease^62^.

### Pathway analysis

Five gene-sets were significantly enriched in broad depression. Of these four were associated with cellular components (where the genes are active) and one was associated with a biological process. The cellular components were all associated with parts of the nervous system (excitatory synapse, neuron spine, post-synapse and dendrite) and demonstrates that genes that are active in these components could be attributing to depression. Excitatory synapses are the site of release for excitatory neurotransmitters and the most common excitatory neurotransmitter, glutamate, has previously been associated with MDD^63^. Neuron spines (or dendritic spines) are extensions from dendrites that act as a primary site for excitatory transmission in the brain. Imbalances of excitation and inhibition have been previously associated with other mental disorders; schizophrenia^64^, Tourette’s syndrome^65^ and autism spectrum disorder^64^. These results indicate that the role of excitatory synapses in the pathology of depression should be further investigated. The biological process, mechanosensory behaviour, refers to behaviour that is prompted from a mechanical stimulus (e.g. physical contact with an object). This indicates that depressive individuals may have a differing behavioural response to mechanosensory stimuli, with higher pain sensitivity and lower pain pressure thresholds found in depression cases^66^.

Our study analysed a large single population-based cohort and replication of the significant genes and variants was not sought. Follow-up analyses should be undertaken in additional cohorts to provide validation of the results obtained in this study. Although useful for studies of this kind, each of the three MDD phenotypes have limitations. None are based on a formal structured diagnostic assessment (such as the Structured Clinical Interview for DSM Axis 1 Disorders interview) and both the broad depression and probable MDD phenotypes are based on self-reported information, which can be subject to recall biases. Broad depression is also likely to be endorsed by a wider range of individuals than traditional depression definitions, including those with internalising disorders other than depression and those with depressive symptoms that would not meet diagnostic criteria for MDD. The ICD-coded MDD phenotype is based on hospital admission records, which can sometimes be incomplete.

### Conclusion

In a large genome-wide association analysis of a broad depression phenotype in UK Biobank, we identified 17 risk variants implicating perturbations of excitatory neurotransmission in depression and high genetic correlations with more comprehensive interview based methods. These findings suggest that a broad depression phenotype may provide a more tractable target for future genetic studies, allowing the inclusion of many more samples. These findings also provide new genetic instruments for the discovery of disease mechanisms, pharmacological treatments and potentially modifiable factors.

## Acknowledgements

This research has been conducted using the UK Biobank Resource – application number 4844. We are grateful to the UK Biobank and all its voluntary participants. The UK Biobank study was conducted under generic approval from the NHS National Research Ethics Service (approval letter dated 17th June 2011, Ref 11/NW/0382). All participants gave full informed written consent.

IJD is supported by the Centre for Cognitive Ageing and Cognitive Epidemiology, which is funded by the Medical Research Council and the Biotechnology and Biological Sciences Research Council (MR/K026992/1). AMMcI, IJD and T-KC acknowledge support from the Wellcome Trust (Wellcome Trust Strategic Award “STratifying Resilience and Depression Longitudinally” (STRADL) Reference 104036/Z/14/Z and the Dr Mortimer and Theresa Sackler Foundation. This investigation represents independent research part-funded by the National Institute for Health Research (NIHR) Biomedical Research Centre at South London and Maudsley NHS Foundation Trust and King’s College London. The views expressed are those of the authors and not necessarily those of the NHS, the NIHR or the Department of Health. DJS supported by Lister Institute Prize Fellowship 2016-2021.

